# Caldas meets Janzen: Thermal regimes of montane plants and implications for global patterns of speciation

**DOI:** 10.1101/2024.05.28.596313

**Authors:** Adriana Sanchez, Ignacio Quintero, Sara Pedraza, Diana Bonilla, Lúcia G. Lohmann, Carlos Daniel Cadena, Felipe Zapata

**Affiliations:** Programa de Biología, Facultad de Ciencias Naturales, Universidad del Rosario, Bogotá, D.C. 111221, Colombia; Institut de Biologie de l’ENS (IBENS), Département de Biologie, École Normale Supérieure, CNRS, INSERM, Université PSL, 75005 Paris, France; Department of Ecology and Evolutionary Biology, University of California, Los Angeles CA 90095, USA; Department of Integrative Biology and Jepson Herbarium, University of California, Berkeley CA 94720, USA; Departamento de Botânica, Instituto de Biociências, Universidade de São Paulo, Rua do Matão 277, Butantã, São Paulo 05508-090, Brazil; Departamento de Ciencias Biológicas, Facultad de Ciencias, Universidad de los Andes, Bogotá, D.C. 111711, Colombia; Center for Tropical Research, Institute of the Environment and Sustainability, University of California, Los Angeles CA 90095, USA

**Keywords:** allopatric, angiosperm, elevation, gradient, latitude, parapatric

## Abstract

The seasonality hypothesis posits that limited seasonal temperature variability in tropical mountains leads to greater climatic zonation along elevation gradients compared to temperate regions. This is predicted to result in narrow thermal tolerances and restricted dispersal for organisms, which may reduce gene flow and increase opportunities for climate-associated parapatric or allopatric speciation in tropical mountains relative to temperate-zone mountains. This hypothesis has been tested in various animal groups but not in plants. We examine the elevational and thermal ranges of pairs of sister species of angiosperms from mountains worldwide. Our findings indicate no significant difference in the breadth and overlap of elevational ranges between tropical and temperate species. However, tropical species have narrower thermal ranges and show greater similarity in these ranges between sister species compared to temperate ones. Such narrow thermal specialization in tropical plants facilitates population divergence and allopatric speciation within thermal zones more than in temperate species.

## Introduction

Understanding the drivers of speciation in tropical mountains–some of Earth’s warmest biodiversity hotspots–is a central question in biogeography, macroecology, and evolutionary biology (Perrigo *et al*. 2020; Rahbek *et al*. 2019a, b). Because geographic variation in climate helps explain differences in global patterns of species diversity (Coelho *et al*. 2023; Kreft & Jetz 2007; Rosenzweig 1995), hypotheses linking differences in climatic variability at lower and higher latitudes with speciation mechanisms could shed light on this open question (Fischer 1960; Ghalambor *et al*. 2006; Quintero & Jetz 2018). In a paper far ahead of its time, Colombian naturalist Francisco José de Caldas noted that climatic variation along elevational and latitudinal gradients influenced the physiology and geographic distribution of organisms (Caldas 1808). Based on his knowledge of astronomy and climatology, Caldas also developed a method to calculate the elevation of mountains and produced profiles of different Andean mountains depicting the distribution ranges of many plant species (González-Orozco *et al*. 2015; Nieto 2006; Vila 2018) (Fig. 1). Likewise, the iconic profile of the Chimborazo volcano made famous by Alexander von Humboldt showing the elevational zonation of plant species (von Humboldt & Bonpland 2010) has long been an inspiration for many ecologists and biogeographers interested in links among biodiversity, climate, and elevation. Notably, such ideas and new insights from field observations formed the core of the seasonality hypothesis (Janzen 1967), which links climate, geography, and physiological performance, and leads to predictions about the distribution of organisms and drivers of speciation in mountain ranges (Ghalambor *et al*. 2006; Sheldon *et al*. 2018).

**Figure 1.**
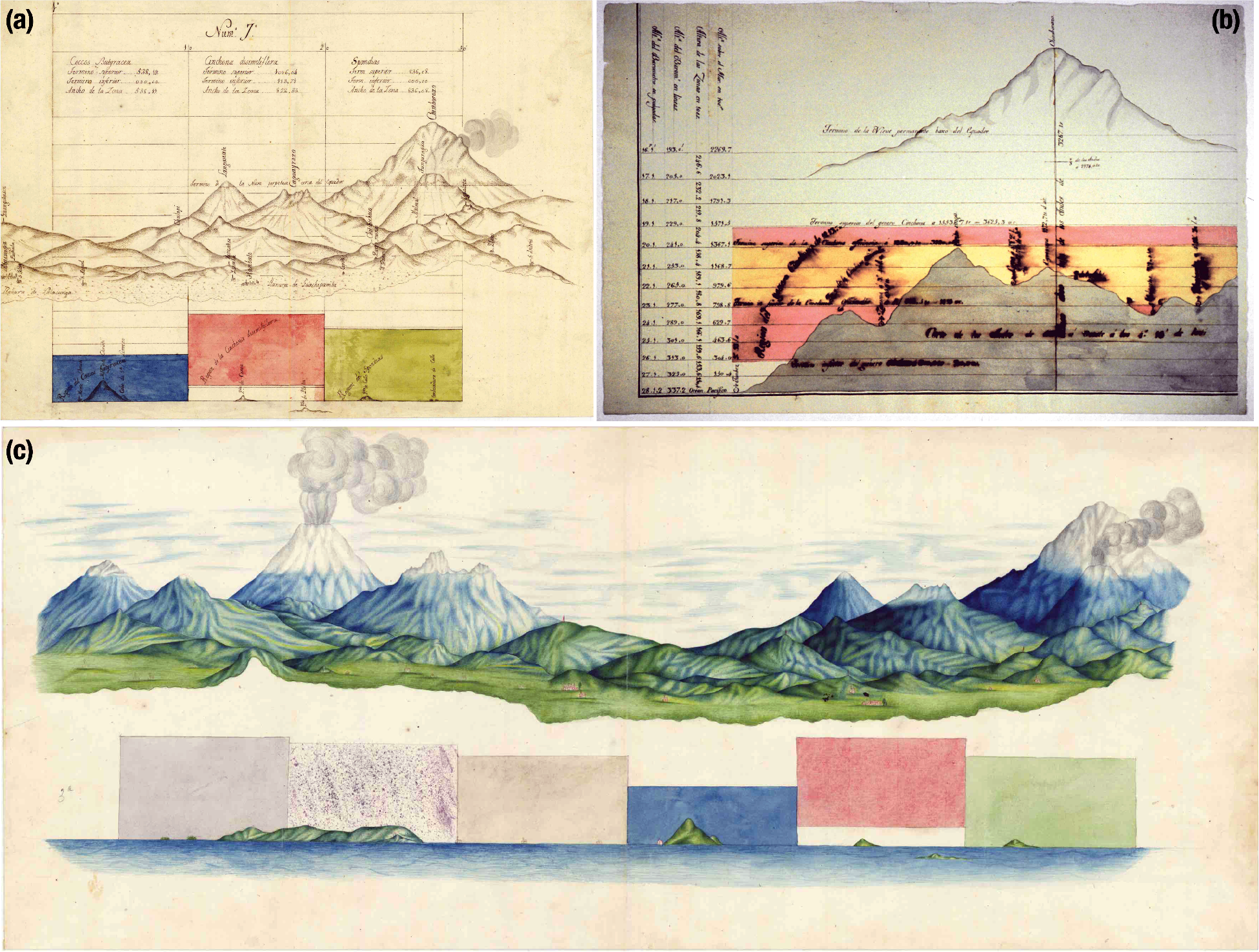
Examples of maps produced by Francisco José de Caldas between 1801 and 1810 illustrating the restricted elevational distributions of tropical plants. (a) Profiles of the Chimborazo volcano on the top right, the Caguyrazo volcano in the center, and the Langanate volcano on the left including designations of zones defined by the dominant plant group, elevation, and three taxon regions described at the bottom of the map: *Spondias* region (in green), *Cinchona dissimiflora* region (in red), and *Cocos butyracca* region (in blue). (b) Elevational profiles of different species of quina plants (*Cinchona*), with that of *Cinchona officinalis* highlighted in yellow. (c) Topographic map of the region between Loja and Quito, Ecuador. All figures from Archivo Cartográfico y de Estudios Geográficos del Centro Geográfico del Ejército, Ministerio de Defensa, Madrid, España -SGE; in (Nieto 2006), reproduced with permission from the author)

Janzeńs (1967) hypothesis states that seasonal variability of climate (i.e., temperature) in tropical mountains is limited relative to that of their temperate counterparts, resulting in greater climatic zonation (i.e. lower climatic overlap) along elevation gradients at lower latitudes.

Consequently, tropical mountain passes –and low-lying valleys– are more likely to act as strong physiological barriers leading to the evolution of narrow thermal tolerances and limiting organismal dispersal up or downslope. Such limited dispersal should result in reduced gene flow, increased local adaptation, and climate-driven geographic isolation in montane tropical populations compared to temperate populations. Over time, limited gene flow could promote population divergence and subsequent speciation, thereby contributing to the build-up of species richness in tropical mountains (Ghalambor *et al*. 2006).

Limited dispersal and narrow thermal specialization of montane tropical organisms can drive speciation via two mechanisms. First, reduced gene flow among populations inhabiting climatically distinct but adjacent elevational zones along tropical mountain slopes may facilitate divergence via parapatric speciation facilitated by climatic niche evolution (Ghalambor *et al*. 2006; Kozak & Wiens 2007). Alternatively, reduced genetic exchange among isolated populations inhabiting similar elevational and climatic zones but in different mountains may increase opportunities for divergence via allopatric speciation facilitated by climatic niche conservatism (Kozak & Wiens 2007; Wiens 2004). If parapatric speciation is the predominant divergence mechanism in tropical mountains, then tropical sister species should inhabit distinct elevational zones, and thus display limited overlap in their thermal ranges. By contrast, if allopatric speciation within thermal zones occurs more commonly in tropical mountains, then tropical sister species should inhabit similar elevational zones (Cadena *et al*. 2012).

Past studies focused on New World vertebrates and invertebrates found strong support for the seasonality hypothesis and prevalent signal for allopatric speciation in the tropics (Cadena *et al*. 2012; Polato *et al*. 2018, but see Kozak & Wiens 2007). However, despite the central role that plants played in inspiring early naturalists to document the remarkable zonation of species along elevation gradients (Caldas 1808; von Humboldt & Bonpland 2010; Fig. 1), the foundational role of plants in determining terrestrial biomes (Donoghue & Edwards 2014; Mucina 2019; Whittaker 1975; Woodward *et al*. 2004), and the abundant opportunities that plants experience to undergo parapatric speciation relative to animals (Bock *et al*. 2023), no studies have comprehensively evaluated the seasonality hypothesis and its connection with drivers of speciation in this group of organisms.

Owing to their sessile nature and reduced set of mitigating behaviors to cope with climatic variability, plants are ideal organisms to study in the context of the seasonality hypothesis because they may experience stronger selection on their physiological tolerances and, thus greater population isolation (Bradshaw 1972; Ghalambor *et al*. 2006; Huey *et al*. 2002). Further, plant performance is tightly linked to temperature via diverse physiological mechanisms (Leigh *et al*. 2012; Michaletz *et al*. 2015). For instance, temperature is one of the most important factors controlling the functional activity of the photosynthetic apparatus (Yordanov 1986). High temperatures can cause permanent decreases in photosynthesis, reduced leaf growth rates, and, eventually, leaf and plant death (Feeley *et al*. 2020; Scafaro *et al*. 2021; Teskey *et al*. 2015). Low temperatures are equally problematic for photosynthesis. Under cold stress conditions, permanent damage to organelles and tissues, loss of turgor, and leaf death may occur (Baxter 2013). Temperature can also influence other key physiological processes such as leaf and flowering phenology as well as seed germination (Crimmins *et al*. 2009;

Rafferty *et al*. 2020; Rosbakh & Poschlod 2015; Vitasse *et al*. 2018). Therefore, the ability of plants to withstand variations in temperature within their environment is critical for plant fitness. Yet, it remains poorly understood whether the environmental tolerances of montane plants are shaped differently by seasonality in tropical versus temperate areas. Moreover, whether and how any such differences might relate to plant speciation mechanisms at different latitudes is not known.

Here, we synthesize geographic, climatic, and phylogenetic data from flowering plants restricted to mountain regions worldwide to test the seasonality hypothesis and its connection to speciation mechanisms. Specifically, we test whether sister species from tropical mountains generally inhabit narrower elevational and thermal ranges relative to temperate sister species. Further, we examine whether patterns of thermal niche overlap between sister species pairs from tropical mountains are consistent with a parapatric or an allopatric speciation model. The former predicts limited thermal overlap between close relatives, whereas the latter predicts conserved thermal niches.

## Material and Methods

### Taxon sampling: geography, climate, and species pairs

To characterize the global elevational and thermal ranges of mountain flowering plants, we constructed a database of georeferenced records for species obtained from the Global Biodiversity Information Facility (GBIF). We restricted all records to montane areas (above 500 m) based on human observations and preserved specimens. Details of additional filters used to download the data and the DOI for the datasets are included in the Supplementary Material. In total, we downloaded 5,743,607 records. Because occurrence data from GBIF can involve geo-location errors, duplicated records, and geo-location uncertainty, we further filtered the data using standard methods (Zizka *et al*. 2019) and custom scripts available in a public GitHub repository at https://github.com/zapata-lab/ms_angios_climatic_zonation. Lastly, to minimize biases in the estimation of elevation and thermal ranges, we discarded species with less than 10 records and filtered elevation outliers for each species (defined as records in the 2.5% tails of the elevational distribution of each species (Cadena *et al*. 2012)). After filtering, our database included 3,767,582 records.

We used the cleaned geo-referenced database to associate all records with temperature layers at 30 s (∼1 km^2^) resolution obtained from the WorldClim v. 2.1 database (Fick & Hijmans 2017). We extracted the maximum temperature of the warmest month (Bio5) and the minimum temperature of the coldest month (Bio6) for each record to characterize the thermal ranges of all species (see below). Additionally, we extracted monthly minimum and maximum temperatures for each record to estimate annual seasonality and the thermal overlap between sister species pairs (see below).

To select species pairs, we used the most comprehensive dated molecular phylogeny of vascular plants available (74,529 tips) (Jin & Qian 2022). This tree reconciles two dated maximum-likelihood phylogenies generated using supermatrices (Smith & Brown 2018; Zanne *et al*. 2013) and standardizes botanical nomenclature according to different databases (Jin & Qian 2022). From this tree, we extracted all pairs of sister species of flowering plants and filtered the cleaned geo-referenced database to include only the species present in such pairs. Because species pairs were identified in the reconstructed tree with the species that survived to the present, it is plausible that for some species pairs the true sister went extinct. The effect of unobserved speciation events, however, decreases towards the present (Nee *et al*. 1994), and we examine any effect of the divergence times between sister species in our subsequent analyses.

We assigned the north temperate region to species with distributions restricted to latitudes higher than 23.437 decimal degrees, the south temperate region to species with distributions restricted to latitudes lower than -23.437 decimal degrees, and the tropical region to species with distributions restricted to latitudes between the north temperate and south temperate regions. Because the focus of our study was on examining differences in elevational and thermal ranges between temperate and tropical species in relation to speciation, we filtered the geo-referenced database to species pairs in which both sister species were restricted to montane areas in the same geographic region. After all filtering steps, the final database included a total of 1,486 species, with 1,012 north temperate species, 284 south temperate species, and 190 tropical species. On average, each species was represented by 242 occurrences, with an average of 235 occurrences for north temperate species, 401 occurrences for south temperate species, and 44 occurrences for tropical species.

### Response variables

We used the elevation information associated with georeferenced records to estimate the elevational range of each species, defined as the difference between the highest and the lowest elevation value across all its records. To estimate the thermal range of each species, we calculated the difference between the maximum value of Bio5 (the maximum temperature of the warmest month) and the minimum value of Bio6 (the minimum temperature of the coldest month) across all records per species (Cadena *et al*. 2012). Because equivalent elevational ranges in tropical and temperate mountains could encompass distinct thermal ranges, we divided the estimated thermal range of each species by the elevation range of each species and used this standardized value as an additional response variable.

We calculated the elevational range overlap between sister species pairs as the degree of overlap (i.e., the lowest of the maximum elevation values for each species minus the highest of the minimum elevation values for each species) divided by the elevational range of the species with the smaller range (Cadena *et al*. 2012; Kozak & Wiens 2007). This value ranges from 0 (no overlap) to 1 (complete overlap). To estimate the overlap in thermal ranges between sister species pairs, we first used the monthly minimum and maximum temperatures to estimate the monthly temperature range experienced by each species by calculating the difference between the mean maximum and the mean minimum temperature across all of its records. Following previous work (Cadena *et al*. 2012; Kozak & Wiens 2007), we then calculated the annual temperature overlap using the formula:

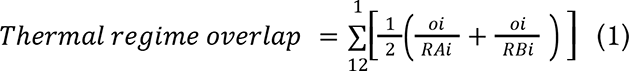

where *R_A_*_i_ and *R_B_*_i_ are the ambient temperature ranges experienced by species *A* and *B* for month *i*, respectively, and *o_i_* is the overlap of *R_A_*_i_ and *R_B_*_i_ (in °C) for month *i*. This annual index of thermal range overlap ranges from 0 to 12, but we rescaled it to a 0 to 1 range before statistical analysis (see below).

Although temperature range overlap obtained from WorldClim data may not actually describe the microclimate that a given species experiences, this approach assumes that there is a relationship between the large-scale variation across the geographical range of a species and its thermal tolerance as expected under the seasonality hypothesis. Other studies have found a positive correlation between macroclimatic data, thermal tolerance of plant species, and latitude (Lancaster & Humphreys 2020; O’sullivan *et al*. 2017).

### Data analyses

We designed a phylogenetic Bayesian modeling framework to test our predictions using the R-INLA package (http://www.r-inla.org), which performs exact posterior parameter inference under the Integrated Nested Laplace Approximation (INLA) (Rue *et al*. 2009). Each regression model was run using the variance-covariance from the phylogenetic tree as the correlation matrix between species while simultaneously estimating lambda to account for phylogenetic non-independence. We specified the default (ambiguous) priors for all parameters, except the hyperparameters of the covariance matrix, for which we used a penalized complexity prior that favors simpler models unless the data support more complexity. To determine whether sister species from tropical mountains generally inhabit narrower elevational, thermal, and standardized ranges relative to sister species from mountains in temperate zones, we performed a generalized linear model for each response variable with an effect of geographical region and the number of records per species. As geographical regions, we compared north-temperate, south-temperate, and tropical regions, as well as temperate (combining both north- and south-temperate) and tropical regions. We included the number of records in the models to account for any effect of sample size on the estimated elevational and thermal ranges. Because the response variables (elevational, thermal, and standardized ranges) are non-negative, we assumed that they are lognormally distributed. We repeated these analyses using quadratic regressions using the mean latitude value per species as a continuous independent variable.

To examine differences in the extent of overlap in elevational and thermal ranges between sister species pairs from tropical and temperate mountains, we designed phylogenetic generalized linear models for each response variable with an effect of geographic regions and divergence times between sister species. In contrast to the first set of analyses above, here the unit of analysis is the species pair, rather than the individual species, which we took into account by estimating a new variance-covariance matrix of the tree with edges extending until the bifurcation events of the sister species. As before, we compared north-temperate, south-temperate, and tropical regions as well as temperate (combining both north- and south-temperate) and tropical regions. We included divergence times between sister species in the models to account for changes in the geographical or thermal ranges of species after speciation, which can influence inferences about the mode of speciation (Losos & Glor 2000) and because the ages of plant sister species may vary with latitude (Igea & Tanentzap 2020). We used the beta distribution for the probability distribution of the response variables (elevational and thermal ranges overlap). We repeated these analyses using quadratic regressions using the mean latitude value per sister species pair as a continuous independent variable. However, because our measures of overlap contained zeros and ones, which are not in the beta distribution, we assumed a censorship value such that a 0 is assumed to be a 0.00001. We checked that this assumption did not change the significance of the relationship with continuous latitude using zero-and-one inflated beta regressions using the *zoib* package for R (Liu & Kong 2015). The scripts used to carry out all statistical analyses are available in a public GitHub repository at https://github.com/zapata-lab/ms_angios_climatic_zonation.

## Results

We found that elevational ranges of tropical flowering plant species were slightly wider than those from temperate mountains when we combined species from the north and south temperate mountains into a single temperate region (Fig. 2a). When we analyzed species from the north and south temperate mountains separately, the south-temperate species showed narrower elevational ranges relative to the species from north-temperate and tropical mountains (Fig. 2b). Together, these results do not support the prediction that species from montane tropical regions occur over narrower elevational ranges than montane temperate species. By contrast, analyses combining species from the north- and south-temperate mountains into a single temperate region showed that species from such latitudes occur over broader temperature ranges than species from tropical mountains (Fig. 2c). When we separated species from the north- and south-temperate mountains, the south-temperate species did not differ from the species in the tropical mountains, and both these groups of species showed narrower thermal ranges compared to the north-temperate species (Fig. 2d). These results support the prediction that species from tropical mountains occur over narrower thermal ranges than species from temperate mountains, particularly when considering mountains in the north temperate region. The distinction between tropical and temperate montane species was notably clearer when we standardized thermal ranges by elevation ranges (Fig 2e). This result persisted when north- and south-temperate mountains were combined into a single temperate region or analyzed independently (Fig. 2f). This finding indicates that species from tropical mountains tend to experience narrower thermal ranges with respect to elevation compared to those in temperate mountains, lending support to the prediction of the climate seasonality hypothesis. The effect of the number of records, while positive and significant, was negligible in all analyses (non-scaled effect < 0.001; Table S1). Results of the quadratic regressions are consistent with the results presented here and can be found in the Supplementary Material (Table S2, Fig. S1)

**Figure 2.**
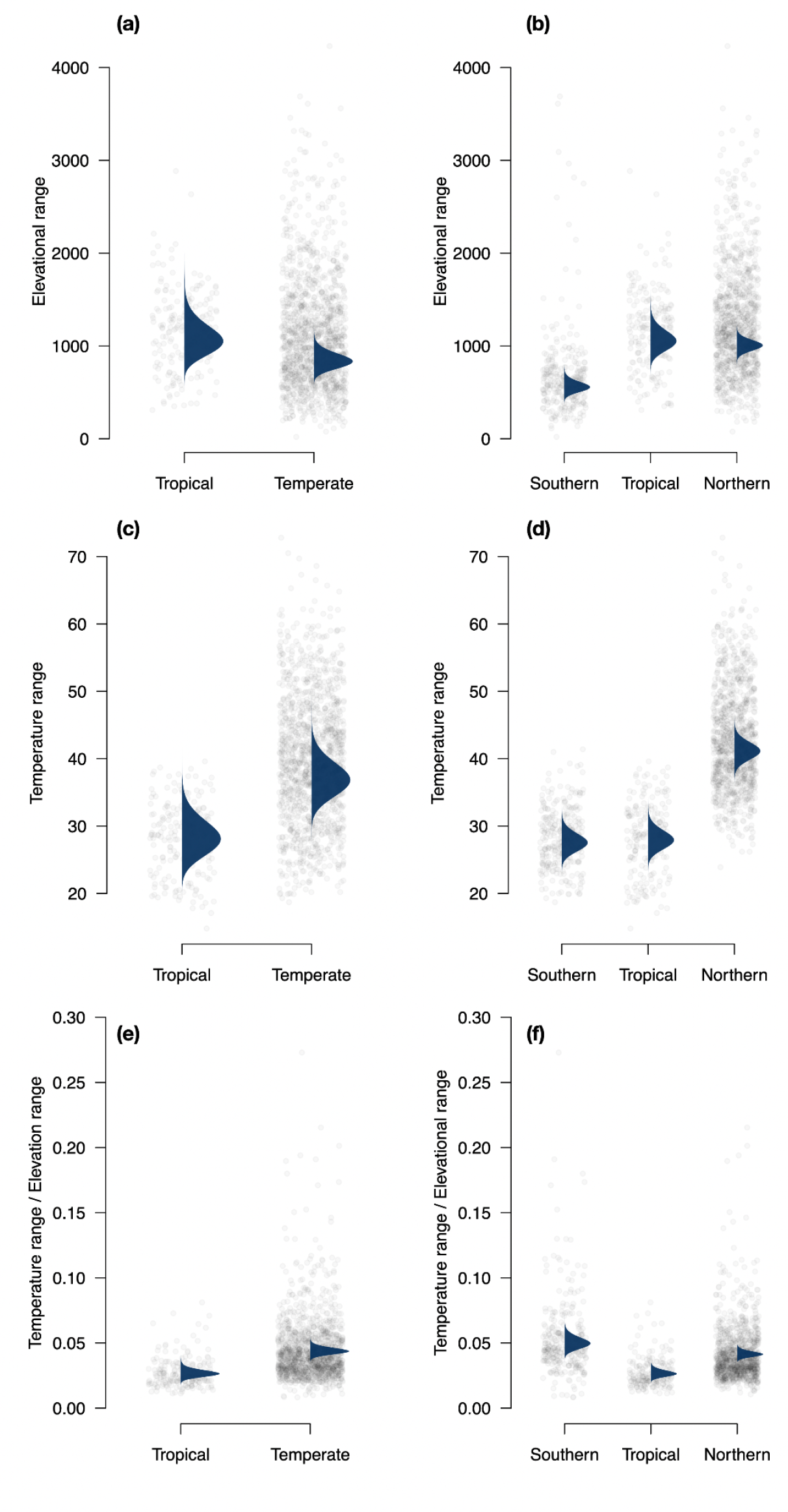
Sister species of flowering plants from tropical mountains worldwide do not occur over narrower elevational ranges than sister species from temperate mountains (a), in particular from the northern hemisphere (b), as evidenced by the overlap in the posterior marginal effect of geographical region. Yet, tropical sister species occur over narrower thermal ranges relative to temperate sister species (c), in particular from the northern hemisphere (d), as evidenced by the non-overlap in the posterior marginals of the geographic region effects. This difference becomes clearer when thermal ranges are standardized by elevation ranges for temperate and tropical species (e) and when temperate species are separated into north- and south-temperate mountains (f). Each panel shows a dotplot of elevation (a, b) and temperature (c, d) ranges, and the standardized temperature by elevation ranges (e, f) between sister species pairs of flowering plants from across geographic regions globally. In (a), (c) and (e), sister species from north and south temperate mountains are combined into a single temperate region. The posterior marginal effects of the geographical region estimated via Bayesian generalized linear models are overlaid in a darker color.

Our analyses showed no differences in the overlap of elevation ranges between sister species pairs from temperate and tropical mountains. These results remained the same when species from the north and south temperate mountains were combined into a single temperate region or analyzed independently (Fig. 3a,b). Conversely, we found a greater overlap in thermal ranges between sister species from tropical mountains relative to sister species from temperate mountains. This result was recovered when we combined sister species from the north and south-temperate mountains into a single temperate region (Fig. 3c). When we analyzed sister species from the north- and south-temperate mountains separately, the overlap in thermal ranges of sister species from the south temperate mountains did not differ from that of the sister species from the tropical mountains, yet both these groups of species pairs showed greater overlap in thermal ranges than sister species pairs from the north temperate mountains (Fig. 3d). In this set of overlap regressions there was a weak, yet significant, negative effect of divergence times in the degree of temperature but not elevation overlap of sister species, reflecting the expected niche conservatism in temperature rather than in elevation per se (Table S3). Results of the quadratic regressions are consistent with the results presented here and can be found in the Supplementary Material (Table S4, Fig. S2). Finally, our zero-and-one-inflated beta regression results confirmed the latitudinal differences in temperature and not in elevation overlap, supporting our INLA results using censorship values (Table S5, Fig. S3).

**Figure 3.**
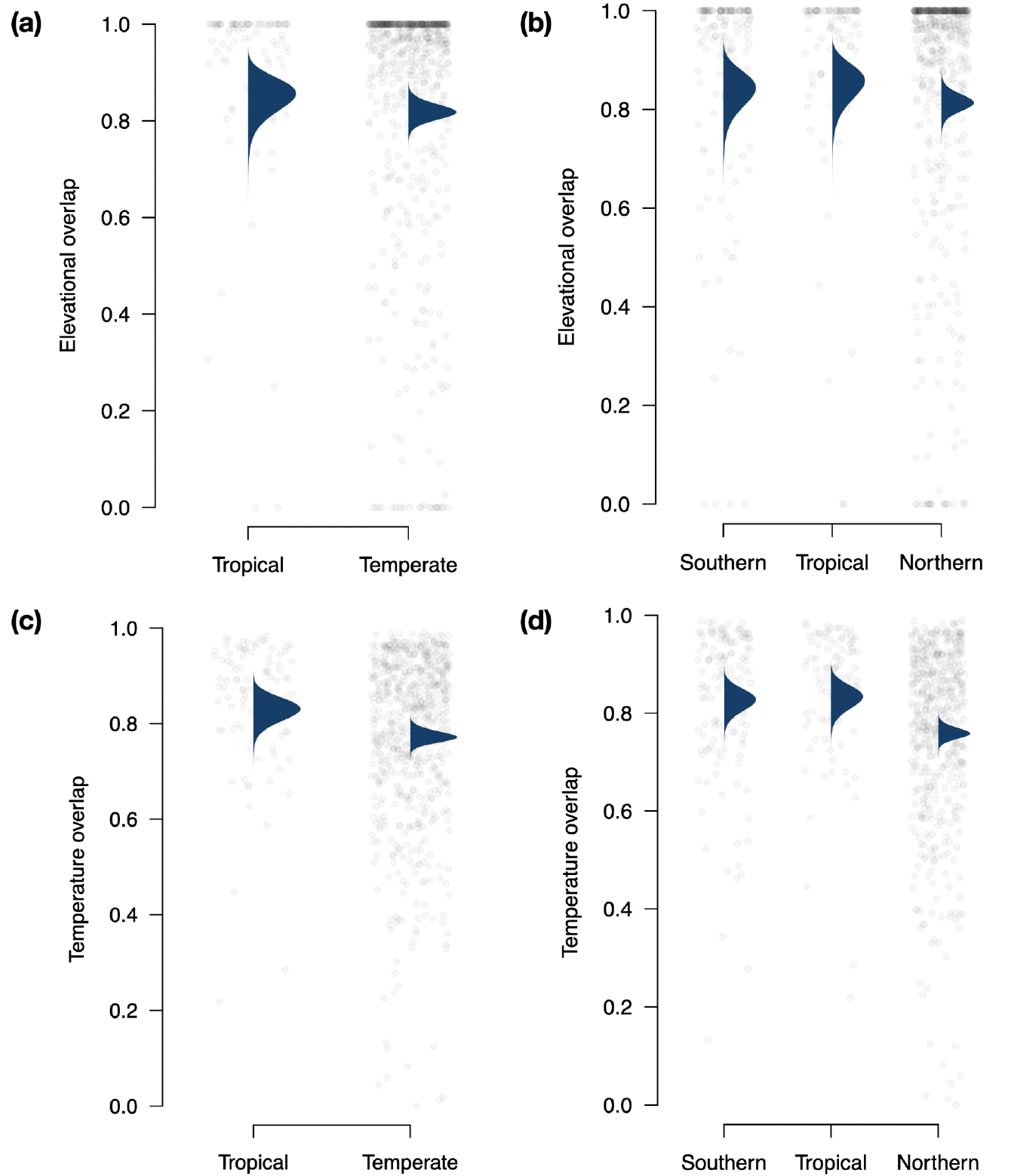
Sister species of flowering plants from tropical mountains worldwide do not differ in the overlap of elevational ranges relative to sister species from temperate mountains (a, b), as evidenced by the overlap in the posterior marginal effect of the geographical region. Yet, tropical sister species show greater overlap in thermal ranges than temperate sister species (c), in particular from the northern hemisphere (d), as evidenced by the minimal overlap in the posterior marginals of the geographic region effects. Each panel shows a dotplot of elevation (a, b) and temperature (c, d) overlap between sister species pairs of flowering plants from across geographic regions globally. In (a) and (c), sister species pairs from north and south temperate mountains are combined into a single temperate region. The posterior marginal effects of the geographical region estimated via Bayesian generalized linear models are overlaid in a darker color.

## Discussion

Using an integrative approach combining geographic, climatic, and phylogenetic information, we revealed that flowering plant species from tropical mountains occur over narrower thermal ranges than species from temperate mountains worldwide despite having similar elevational ranges (Fig 2).

This result is consistent with elements of the seasonality hypothesis (Ghalambor *et al*. 2006; Janzen 1967; Sheldon *et al*. 2018). Further, we showed that the thermal ranges of montane tropical plants overlap more extensively than the thermal ranges of species from temperate montane regions (Fig. 3). Taken together, these findings suggest that the narrow thermal specialization of tropical flowering plants may offer greater opportunities for population divergence via allopatric speciation within thermal zones compared to species from the temperate zone.

A recent study comparing the thermal tolerance breadth of plant species from a few localities in the mountains of Bolivia, Ecuador, USA, and Austria reported that species from tropical regions did not have narrower thermal tolerances relative to temperate-zone species (Sklenář *et al*. 2023). As a result, these authors concluded that the seasonality hypothesis does not hold true for plant species, and therefore that tropical mountain passes may not represent significant physiological barriers for plants. These findings seemingly contradict the findings of our study. However, a closer inspection of the localities included in the study by Sklenář *et al*. (2023) reveals that the tropics were represented only by plants from the highest elevation habitats above 4000 m (i.e., evironments above the treeline including páramos, punas, and jalcas). Previous studies have shown that high-elevation tropical habitats have a broad daily variability in temperature, leading to the “summer every day and winter every night” phenomenon (Hedberg 1964; Vuilleumier & Monasterio 1986). This means that daily temperature fluctuations in alpine tropical montane habitats mirror the seasonal temperature oscillations of temperate mountains (Chan *et al*. 2016; Ghalambor *et al*. 2006). Consequently, the breadth of the thermal ranges experienced by tropical species is expected to increase with elevation, with species at the highest summits operating effectively as temperate-zone species, as has been demonstrated for some animals (Ghalambor *et al*. 2006; Gill *et al*. 2016; McCain 2009; Polato *et al*. 2018). A recent analysis of woody plants along a ca. 3,500 m elevation gradient in Bolivia is consistent with this prediction (Montaño-Centellas *et al*. 2023). This study found a strong positive relationship between temperature variability and the elevational range size of approximately 2,300 plant species. However, another study limited to high-elevation summits (>3,000 m) along a latitudinal gradient in the Andes (8°N - 26°S) found no relationship between thermal tolerance breadth and elevation, but detected a monotonic increase with latitude (Cuesta *et al*. 2020). In contrast to the mixed evidence from these previous findings, our results, encompassing tropical mountains globally and along full elevational gradients, thereby capturing both high- and low-elevation species experiencing more and less variability in temperature, strongly support the seasonality hypothesis. Notably, we show that the tropical and temperate species differ in the thermal range standardized by elevation range, with tropical species experiencing a narrower range of temperatures per elevation range with respect to temperate species (Fig. 2). This implies a greater climatic zonation along elevation in tropical mountains and possibly more climate-associated speciation in the tropics relative to the temperate zones. Future work, however, is needed for a more detailed assessment of how plant thermal tolerance breadth varies across lineages, along elevational gradients, and among mountain regions worldwide (Perez *et al*. 2016).

Our finding of a higher overlap in the thermal regimes of sister species in tropical mountains indicates that thermal niches in tropical plants are more conserved than in temperate-zone species (Fig. 3). The relatively high overlap in thermal regimes between sister species in tropical mountains is consistent with the hypothesis that allopatric speciation is the dominant mode of speciation in the region. Neontological studies in different plant clades and geographic regions have documented patterns of diversification consistent with this idea, revealing that geographical isolation is the prominent driver of plant speciation in tropical mountains (Brochmann *et al*. 2022; Diazgranados & Barber 2017; Vargas *et al*. 2023). In addition, macroecological and macroevolutionary analyses suggest this may be a common phenomenon across the tree of life (Crisp *et al*. 2009; Donoghue 2008).

Notably, a large-scale analysis of vertebrate taxa in the Americas revealed support for the same pattern of higher conservation in thermal regimes for tropical sister species, suggesting that the greater stability of temperature regimes along tropical mountain slopes increases opportunities for isolation and allopatric speciation in animals (Cadena *et al*. 2012). Also, phylogenetic analyses consistently reveal that species replacing each other along tropical elevation gradients are not each other’s closest relatives and that sister taxa typically occur at similar elevations in different mountains, revealing that thermal niches are largely conserved during speciation (Cadena & Céspedes 2020; Linck *et al*. 2021). Thus, a strong pattern is emerging for a generalized geographic speciation mode in montane tropical taxa across organisms with disparate reproductive traits, life histories, and dispersal abilities. More broadly, our study suggests that climatic niche evolution operates similarly in plants and animals despite the marked differences in their biology (Liu *et al*. 2020).

On the other hand, high overlap in the thermal regimes of tropical sister species does not discard alternative speciation modes driven by ecological factors unrelated to the thermal niche. For instance, if plant speciation is driven by changes in ploidy level (i.e., polyploidization, hybridization) or switches in biotic interactions that influence the reproductive success and fate of populations (e.g., pollinator shifts, allochrony in flowering time), populations might differentiate and speciate in sympatry. While it is indeed plausible that these factors may be common in plants and can lead to a marked prevalence of sympatric speciation in a few plant taxa (Bock *et al*. 2023; Skeels & Cardillo 2019), it is difficult to envision how such speciation mode might be more likely to happen only in tropical mountains across many species pairs of flowering plants sampled from across the angiosperm phylogeny. We argue instead that our results reflect the fact that allopatric speciation is the dominant speciation mode in plants (and likely animals) in the mountains closer to the equator.

In contrast to the predictions of the seasonality hypothesis, we did not find that sister species from tropical mountains had narrower elevational ranges than sister species from temperate-zone mountains (Fig. 2a,b). We also found that sister species pairs from temperate mountains in the southern hemisphere experience temperate ranges and thermal overlaps more similar to those of tropical sister pairs than their north temperate counterparts (Figs. 2, 3). These north-south differences are consistent with disparities in the climates of the two hemispheres (Chown *et al*. 2004).

Temperatures in the southern hemisphere are less extreme than those in the north, especially in the winter, exposing organisms to milder yearly temperatures on average. This is in part related to the geometry of continental land masses and the predominance of a continental climate in the north relative to an oceanic climate in the south. Thus, the limited seasonal variability temperature in the southern hemisphere can result in a greater climatic zonation along elevation gradients in south temperate mountains, akin to that in tropical mountains (Janzen 1967). Our results are consistent with this hypothesis and reveal that the thermal niches of south temperate sister species tend to be narrower and more evolutionarily conserved than those of north temperate sister species. As a result, we suggest that plant populations in southern hemisphere mountains generally experience greater opportunities for isolation and allopatric speciation, similar to plant populations in tropical mountains. More generally, our results are consistent with the idea that the tropical vs. temperate distinction between montane areas established by Janzen (Janzen 1967) may not fully capture the way in which temperature varies with elevation. Beyond latitude, other factors influence thermal overlap across elevations (Zuloaga & Kerr 2017) and this needs to be better characterized and incorporated into models describing speciation in relation to climatic variation in mountains across the globe. For example, the velocity of climate change along elevation gradients (i.e., the velocity of isotherm shifts) can determine the breadth of species elevation ranges and whether species may track climate warming via range shifts. While the velocity of isothermal shifts along elevation varies drastically worldwide, the highest vertical velocities are concentrated at low elevations, mostly in the north-temperate mountains (Chan *et al*. 2024). Establishing whether and how such shifts may influence speciation mechanisms remains key for assessing the large-scale effect of temperature variability in mountain speciation and its consequences in the context of global climate change.

Our inferences assume that temperature variation measured at a macroclimatic scale can accurately delineate plant species ranges and capture meaningful physiological properties about organismal performance. Certainly, temperature is a dominant factor governing multiple processes related to plant fitness (Taiz *et al*. 2022) and broad-scale studies show that plant thermal tolerance decreases with latitude (O’sullivan *et al*. 2017) and elevation in the tropics (Feeley *et al*. 2020). Thus, based on the available evidence, our results align with plant thermal physiology and macroecological trends. Although additional environmental factors, including soil properties, drought, radiation, biotic interactions, and land cover can influence plant species ranges and physiology (Chauvier *et al*. 2021; Maharjan *et al*. 2022; Pellissier *et al*. 2013), it is not well understood how such factors vary along elevation gradients globally and therefore relate to speciation modes in montane plants. Our analyses also use macroclimatic data and yearly averages to estimate thermal ranges and thermal overlap. Yet, if the actual operational thermal environment experienced by plants is limited temporally or spatially, our results may overestimate the realized thermal ranges. For instance, if the short-lived annual habit and deciduousness are more common in places with high seasonality and limited to certain periods, temperature will only be a critical factor at a particular point in time rather than continuously year-round. Similarly, mounting evidence shows that the temperature experienced by the organs involved in plant thermal eco-physiology can vary at finer spatial scales, including microhabitats and forest strata (Klinges & Scheffers 2020; Vinod *et al*. 2023). Therefore, future studies should investigate to what extent variations in microhabitat preferences, vertical forest profiles, life-history strategies, leaf morphologies, and thermoregulatory mechanisms between sister species, both in temperate and tropical mountains, influence their thermal niches (Pedraza 2024).

In sum, this study presents evidence of narrower and highly conserved thermal niches in plant sister species from tropical mountains relative to sister species from temperate mountains, particularly from the northern hemisphere. The observed pattern of thermal niche conservatism suggests allopatric speciation as a dominant mode of speciation for plants in tropical mountains and possibly in mountains from the southern hemisphere where seasonal climatic variation is less extreme than in the northern hemisphere. However, exactly why species from the tropics and south temperate regions exhibit greater conservatism in their thermal niche will require further study. For instance, comparative genomic and physiological analyses can illuminate the mechanisms of local thermal adaptation, which should be more prevalent in tropical and south temperate plant species than in species from north temperate mountains. Furthermore, detailed comparative population genetic surveys assessing the extent of gene flow along elevation and across latitude will be necessary to quantify the strength of mountain passes in driving population isolation and plant speciation. The asymmetry in the pattern of climatic niche conservatism reported here also suggests that plant species from tropical mountains, and likely south temperate mountains, can track their preferred climates across geography but lack the ability to adapt to new climatic conditions, making them particularly vulnerable to rapid global warming (Feeley *et al*. 2023; Linck *et al*. 2021).

## Acknowledgments

We are grateful to Michael Landis for his feedback on this work.

## Supplementary Methods

We downloaded georeferenced records from the Global Biodiversity Information Facility (GBIF) using the following filters. An explanation of the filter options can be found on the GBIF website https://www.gbif.org/

*Basis of record*: Human observation, preserved species

*Elevation*: 500 - 9,999 m

*Has coordinate*: true

*Has geospatial issue*s: false

*Issues and flag*s: coordinate rounded, Geodetic datum assumed WGS84, Coordinate reprojected, Country derived from coordinates, Occurrence status inferred from individual count, Ambiguous institution, Ambiguous collection, Institution match none, Collection match none, Institution match fuzzy, Collection match fuzzy, Institution collection mismatch, Different owner institution.

*Occurrence status*: present

*Scientific name*: Liliopsisa, Magnoliopsida *Year*: between start of 1946 and end of 2022. **DOI of each dataset**

DOI Liliopsida: 10.15468/dl.8etn3b DOI Magnoliopsida: 10.15468/dl.afyzay

## Supplementary Tables and Figures

**Table S1.**
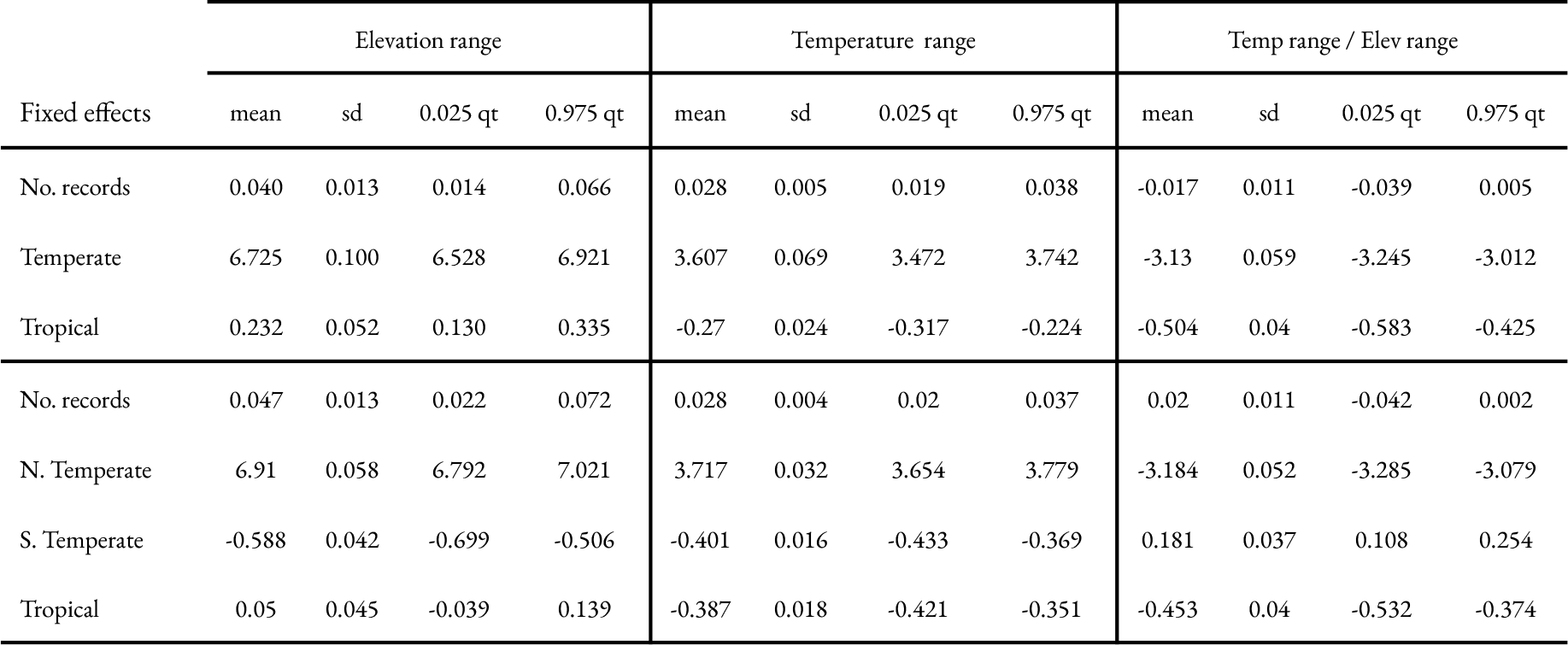
Posterior marginal effects for the Bayesian generalized linear models analyzing the relationship between the geographical region and the number of records per species, with the elevation range, the temperature range, and the temperature range standardized by elevation range. The models include the mean (mean), standard deviation (sd), and 95% credible intervals for the coefficients of the fixed effects. The credible intervals provide a range within which the true parameter values lie with 95% probability, offering a measure of the uncertainty associated with each estimate.

**Table S2.** Posterior marginal effects for the Bayesian quadratic regressions analyzing the relationship between the mean latitude value per species and the number of records per species, and elevation range, the temperature range, with the temperature range standardized by elevation range. The model includes the mean (mean), standard deviation (sd), and 95% credible intervals for the coefficients of the predictor variable. The credible intervals provide a range within which the true parameter values lie with 95% probability, offering a measure of the uncertainty associated with each estimate.

| Fixed effects | Elevation range |  |  |  | Temperature range |  |  |  | Temp range / Elev range |  |  |  |
| --- | --- | --- | --- | --- | --- | --- | --- | --- | --- | --- | --- | --- |
|  | mean | sd | 0.025 qt | 0.975 qt | mean | sd | 0.025 qt | 0.975 qt | mean | sd | 0.025 qt | 0.975 qt |
| Intercept | 6.895 | 0.058 | 6.78 | 7.007 | 3.527 | 0.044 | 3.44 | 3.614 | -3.363 | 0.069 | -3.498 | -3.226 |
| No. records | 0.043 | 0.013 | 0.018 | 0.068 | 0.028 | 0.005 | 0.019 | 0.037 | -0.018 | 0.012 | -0.041 | 0.006 |
| Latitude | 0.149 | 0.03 | 0.089 | 0.280 | 0.208 | 0.013 | 0.184 | 0.233 | 0.089 | 0.029 | 0.032 | 0.146 |
| Latitude<sup>2</sup> | -0.066 | 0.021 | -0.108 | -0.024 | 0.034 | 0.009 | 0.016 | 0.051 | 0.117 | 0.021 | 0.076 | 0.157 |

**Table S3.** Posterior marginal effects for the Bayesian generalized linear models analyzing the relationship between the geographical region, the divergence time between species pairs, and the minimum number of records per species pair, with the overlap in elevation and temperature ranges. The models include the mean (mean), standard deviation (sd), and 95% credible intervals for the coefficients of the fixed effects. The credible intervals provide a range within which the true parameter values lie with 95% probability, offering a measure of the uncertainty associated with each estimate.

| Fixed effects | Elevation overlap |  |  |  | Temperature overlap |  |  |  |
| --- | --- | --- | --- | --- | --- | --- | --- | --- |
|  | mean | sd | 0.025 qt | 0.975 qt | mean | sd | 0.025 qt | 0.975 qt |
| No. records | 0.028 | 0.053 | -0.077 | 0.132 | 0.089 | 0.041 | 0.008 | 0.169 |
| Pair age | -0.018 | 0.014 | -0.046 | 0.009 | -0.022 | 0.009 | -0.04 | -0.004 |
| Temperate | 1.520 | 0.0123 | 1.29 | 1.78 | 1.229 | 0.073 | 1.094 | 1.382 |
| Tropical | 0.273 | 0.168 | -0.056 | 0.602 | 0.366 | 0.104 | 0.161 | 0.570 |
| No. records | 0.023 | 0.053 | -0.081 | 0.127 | 0.074 | 0.039 | -0.003 | 0.151 |
| Pair age | -0.021 | 0.014 | -0.049 | 0.006 | -0.028 | 0.009 | -0.045 | -0.01 |
| N. Temperate | 1.486 | 0.131 | 1.237 | 1.759 | 1.152 | 0.068 | 1.026 | 1.294 |
| S. Temperate | 0.209 | 0.153 | -0.087 | 0.513 | 0.420 | 0.092 | 0.240 | 0.6 |
| Tropical | 0.318 | 0.172 | -0.019 | 0.656 | 0.463 | 0.105 | 0.258 | 0.669 |

**Table S4.** Posterior marginal effects for the Bayesian quadratic regressions analyzing the relationship between the mean latitude value per species, the divergence time between species pairs, and the minimum number of records per species pair, with the overlap in elevation and temperature ranges. The model includes the mean (mean), standard deviation (sd), and 95% credible intervals for the coefficients of the predictor variable. The credible intervals provide a range within which the true parameter values lie with 95% probability, offering a measure of the uncertainty associated with each estimate.

| Fixed effects | Elevation overlap |  |  |  | Temperature overlap |  |  |  |
| --- | --- | --- | --- | --- | --- | --- | --- | --- |
|  | mean | sd | 0.025 qt | 0.975 qt | mean | sd | 0.025 qt | 0.975 qt |
| Intercept | 1.652 | 0.159 | 1.352 | 1.98 | 1.564 | 0.085 | 1.402 | 1.735 |
| No. records | 0.056 | 0.054 | -0.080 | 0.132 | 0.112 | 0.043 | 0.028 | 0.196 |
| Pair age | -0.021 | 0.014 | -0.049 | 0.007 | -0.028 | 0.009 | -0.045 | -0.011 |
| Latitude | -0.170 | 0.122 | -0.411 | 0.067 | -0.509 | 0.073 | -0.651 | -0.366 |
| Latitude<sup>2</sup> | -0.077 | 0.089 | -0.252 | 0.097 | -0.260 | 0.054 | -0.365 | -0.154 |

**Table S5.**
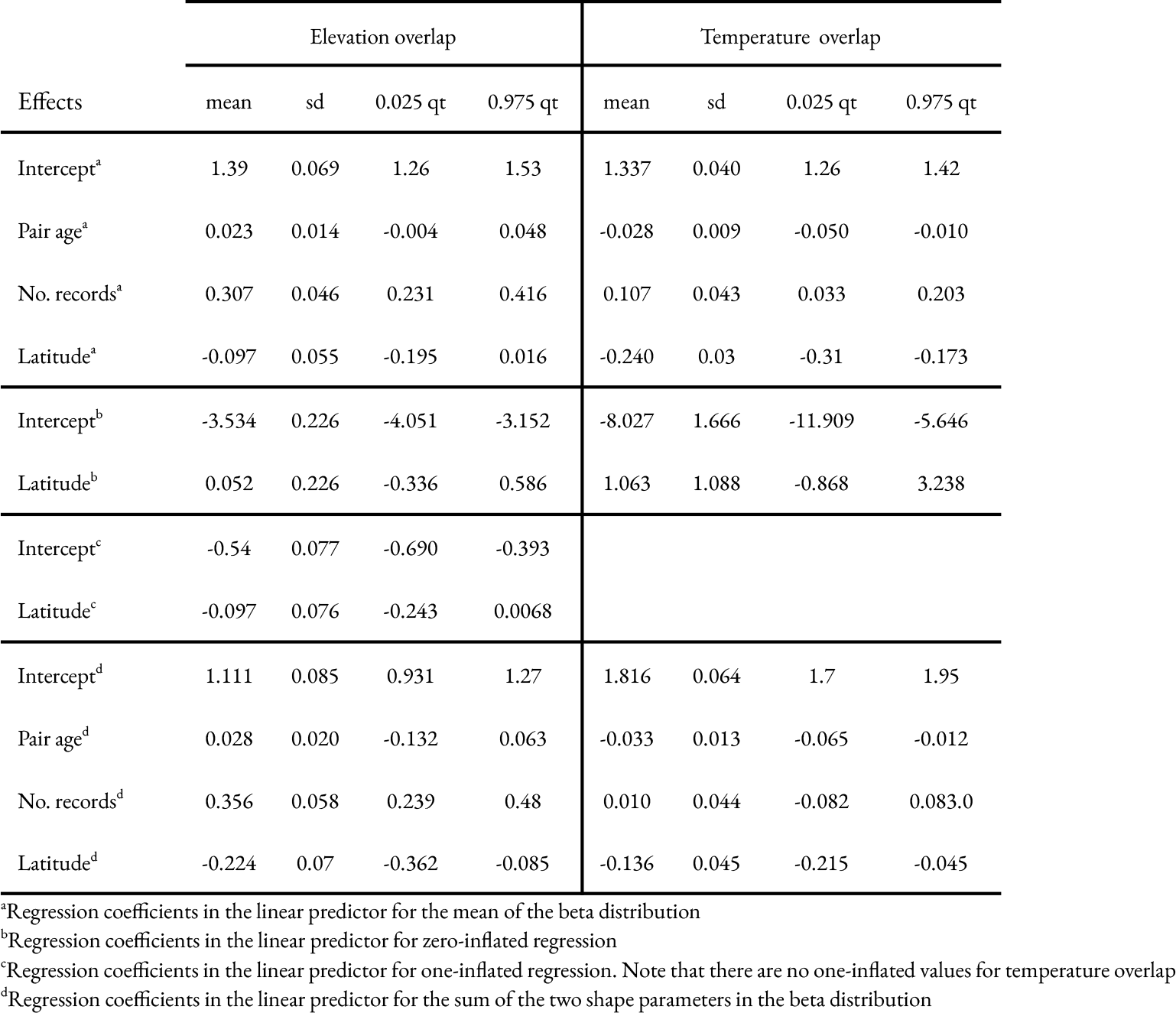
Bayesian posterior inferences of the joint zero-and-one inflated beta regression model parameters. The models include the mean (mean), standard deviation (sd), and 95% credible intervals for the coefficients of the predictor variable. The credible intervals provide a range within which the true parameter values lie with 95% probability, offering a measure of the uncertainty associated with each estimate.

**Figure S1.**
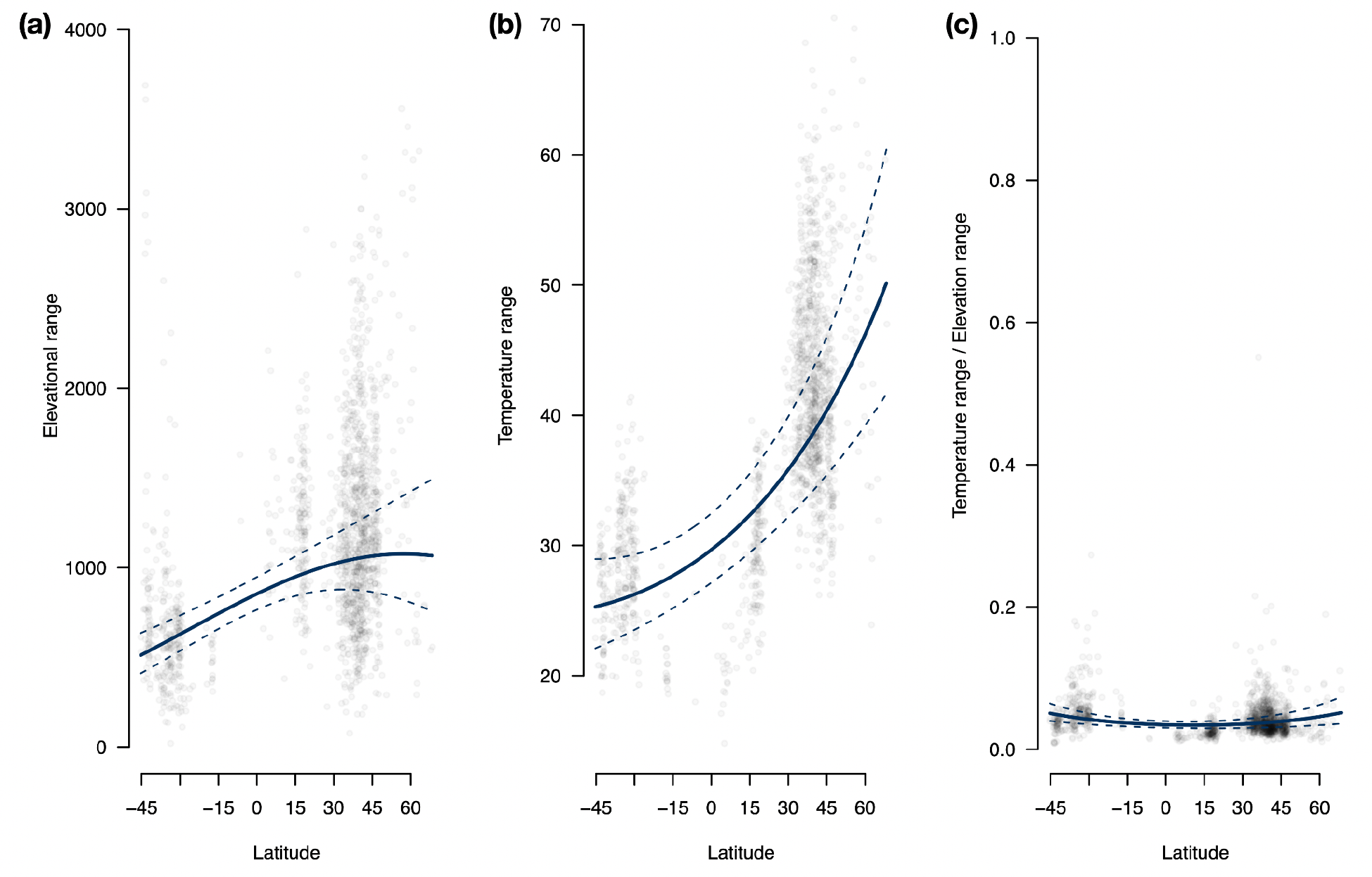
Sister species of flowering plants from tropical mountains worldwide do not occur over narrower elevational ranges than sister species from temperate mountains, in particular from the northern hemisphere (a). Yet, tropical sister species occur over narrower thermal ranges relative to temperate sister species, in particular from the northern hemisphere (b). This difference becomes clearer when thermal ranges are standardized by elevation ranges (c). Each panel shows a dotplot of elevation (a), temperature (b), and the standardized temperature by elevation ranges (c) between sister species pairs of flowering plants from across geographic regions globally. The posterior marginal effects of latitude (continuous line) and 95% credibility intervals (dashed lines) were estimated via Bayesian quadratic regressions.

**Figure S2.**
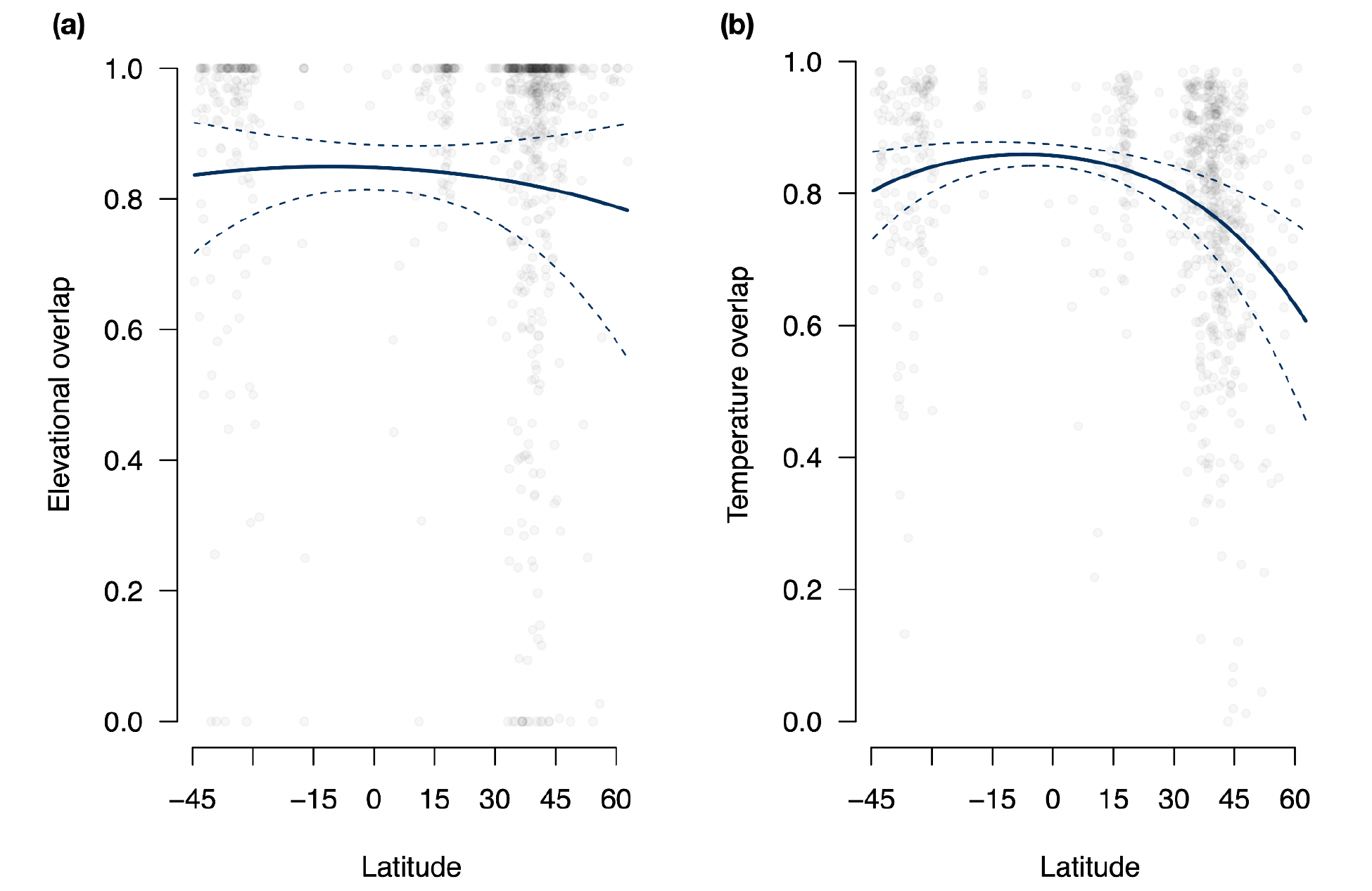
Sister species of flowering plants from tropical mountains worldwide do not differ in the overlap of elevational ranges relative to sister species from temperate mountains (a). Yet, tropical sister species show greater overlap in thermal ranges than temperate sister species (b). Each panel shows a dotplot of elevation (a) and temperature (b) overlap between sister species pairs of flowering plants from across geographic regions globally. The posterior marginal effects of latitude (continuous line) and 95% credibility intervals (dashed lines) were estimated via Bayesian quadratic regressions.

**Figure S3.**
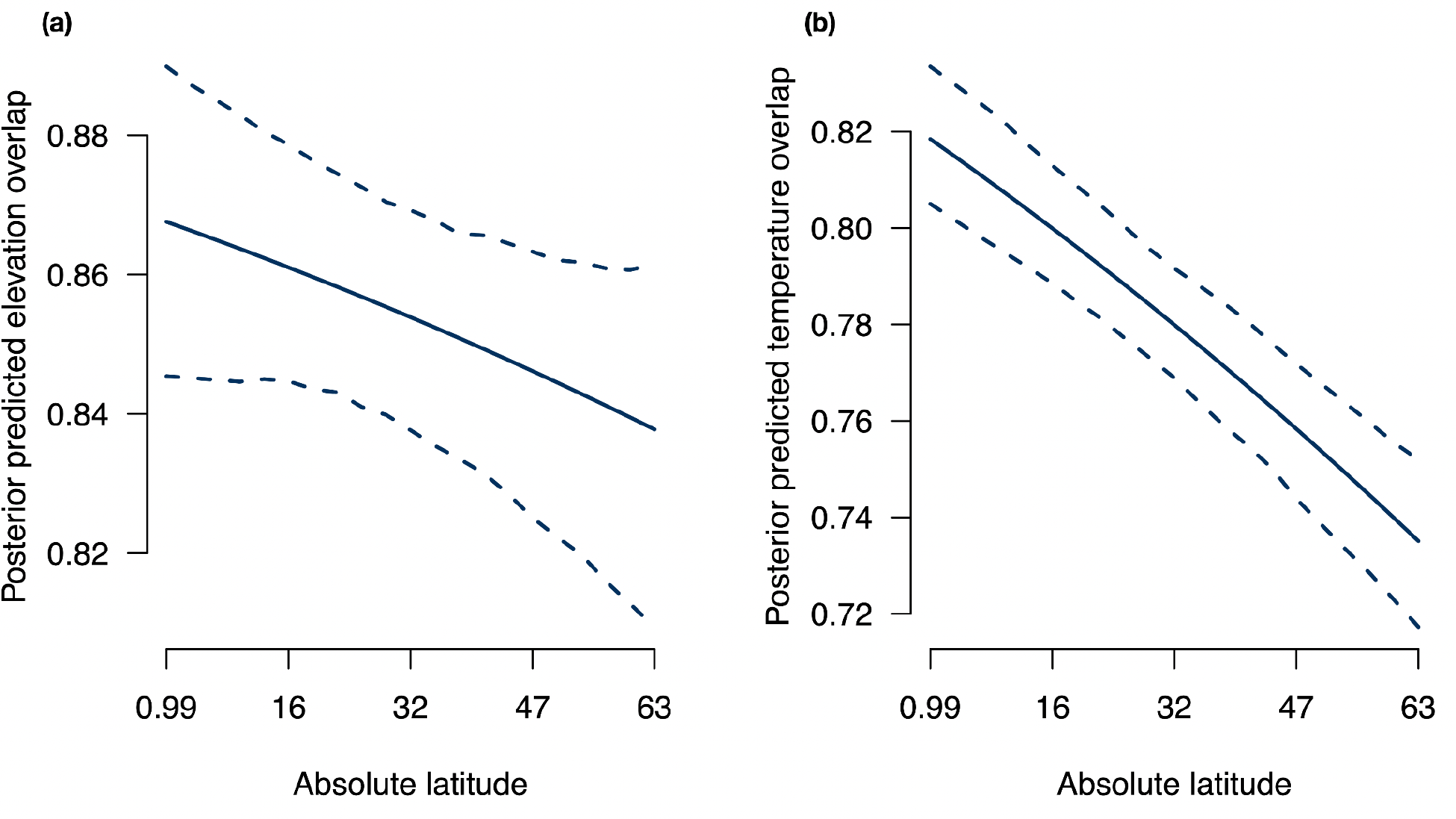
Posterior predictive distribution of the effect of latitude on elevation (a) and thermal (b) overlap while maintaining the number of records per species pair and the divergence time between species pairs constant. The effect of latitude is only significant for temperature overlap (b).

